# PEPE: Scalable extraction of multi-modal protein language model representations

**DOI:** 10.1101/2025.10.13.680902

**Authors:** Jahn Zhong, Niccolò Cardente, Geir Kjetil Sandve, Habib Bashour, Maria Francesca Abbate, Victor Greiff

## Abstract

**Motivation:** Protein language models (PLMs) capture intricate amino-acid dependencies, producing embeddings that encode rich structural, functional, and evolutionary information. Despite their potential, current extraction workflows rely on arbitrary choices, with respect to embedding layer, pooling, and padding, that frequently yield suboptimal representations for feature extraction and downstream analyses. Large-scale embedding generation is further limited by inefficiencies in computation and memory: (i) accumulating all model outputs in memory before writing to disk causes severe bottlenecks, and (ii) repeatedly embedding identical sequences to extract different modes introduces redundant computation and drastically reduces throughput and scalability.

**Results:** We introduce PEPE (Parallel Extraction for Protein Embeddings), a command-line tool and Python library that enables efficient, high-throughput, and multimodal extraction from protein language models. PEPE’s parallelized and streaming-based architecture achieves runtimes several orders of magnitude faster than sequential approaches. Unlike conventional methods—whose peak memory usage scales linearly with output size and fails when memory capacity is exceeded—PEPE maintains stable, low memory consumption, enabling multimodal embedding extraction even beyond available RAM. PEPE supports a wide range of state-of-the-art and custom PLMs through a simple, flexible interface. By combining scalability, robustness, and ease of use, PEPE allows researchers to generate massive, information-rich embedding datasets efficiently, and facilitate the discovery of optimal representations for structural, functional, and evolutionary downstream tasks. By streamlining the generation of diverse embedding configurations, PEPE provides researchers with the necessary data to identify high-performing latent states for specific biological contexts without requiring additional computational resources.

**Availability and Implementation:** PEPE is a command-line tool written in Python and published under MIT license. The source code and documentation are available at https://github.com/csi-greifflab/pepe-cli. PEPE is also available for installation from PyPI under https://pypi.org/project/pepe-cli and deposited on Zenodo at https://zenodo.org/records/15912054.

## 1. Introduction

Recent advances in protein language models (PLMs) have enhanced our ability to learn and interpret the underlying patterns of protein sequences, enabling breakthroughs in computational tasks such as structure prediction ^1–3^, function annotation ^4^, and de novo protein design ^5,6^. Central to these developments is the adoption of transformer-based ^7^, PLMs, such as ESM-2 ^2^, AntiBERTa2-CSSP ^3^, ProtTrans ^8^ or xTrimoPGLM ^9^. Typically, PLMs are pre-trained on large and diverse training datasets using self-supervised masked language modeling tasks ^10,11^, with the aim of learning generalizable feature representations (embeddings) of protein sequences. Pre-trained models are used for transfer learning mainly by: (1) fine-tuning for new tasks with relatively little labeled data ^8,10,12–15^, or (2) using their embeddings as input features for downstream models ^8,10,12–15 14,16–19^.

In this work, we use the term embedding to refer to high-dimensional numerical representations extracted from outputs of PLMs. We call different representations of the same sequence “embedding modes” (i.e., different layers or pooling methods). As each embedding layer modifies the outputs of the previous layer and passes them to the next, embeddings from the final layer are considered the most informative and are often the chosen embedding mode for transfer learning tasks ^20–22^. However, the relevance of deeper layers varies by task, and in some cases intermediate layers could be more informative, especially for rare sequences, such as those of adaptive immune receptors (AIRs) ^20–25^.

In the absence of general “best practices”, the optimal embedding mode for a given transfer learning task may be determined by benchmarking multi-modal embedding datasets that contain multiple embedding modes of each protein sequence. While the design and execution of such benchmarks have become more accessible through modern platforms like ImmuneML^26^, generating the underlying datasets remains challenging. Although state-of-the-art (SOTA) tools, such as PLMFit ^27^ or Bio Embeddings ^28^, are theoretically able to extract many embedding modes of a sequence, in practice, they use a sequential process, which limits scalability due to constraints in memory capacity and throughput (see Figure 1A). The sequential process begins by tokenizing the input protein sequences and dividing them into batches. Then, each batch of tokens is embedded and temporarily stored in memory. Finally, after the outputs of all batches have been stored, they are written to a storage file (extraction). All steps are then repeated for each embedding mode to be extracted, resulting in significant computational overhead. To address these limitations, we designed PEPE (**P**arallel **E**xtraction for **P**rotein **E**mbeddings), where, concurrently with the computation of embeddings, multiple embedding modes can be extracted from a single forward pass through the model.

**Figure 1:**
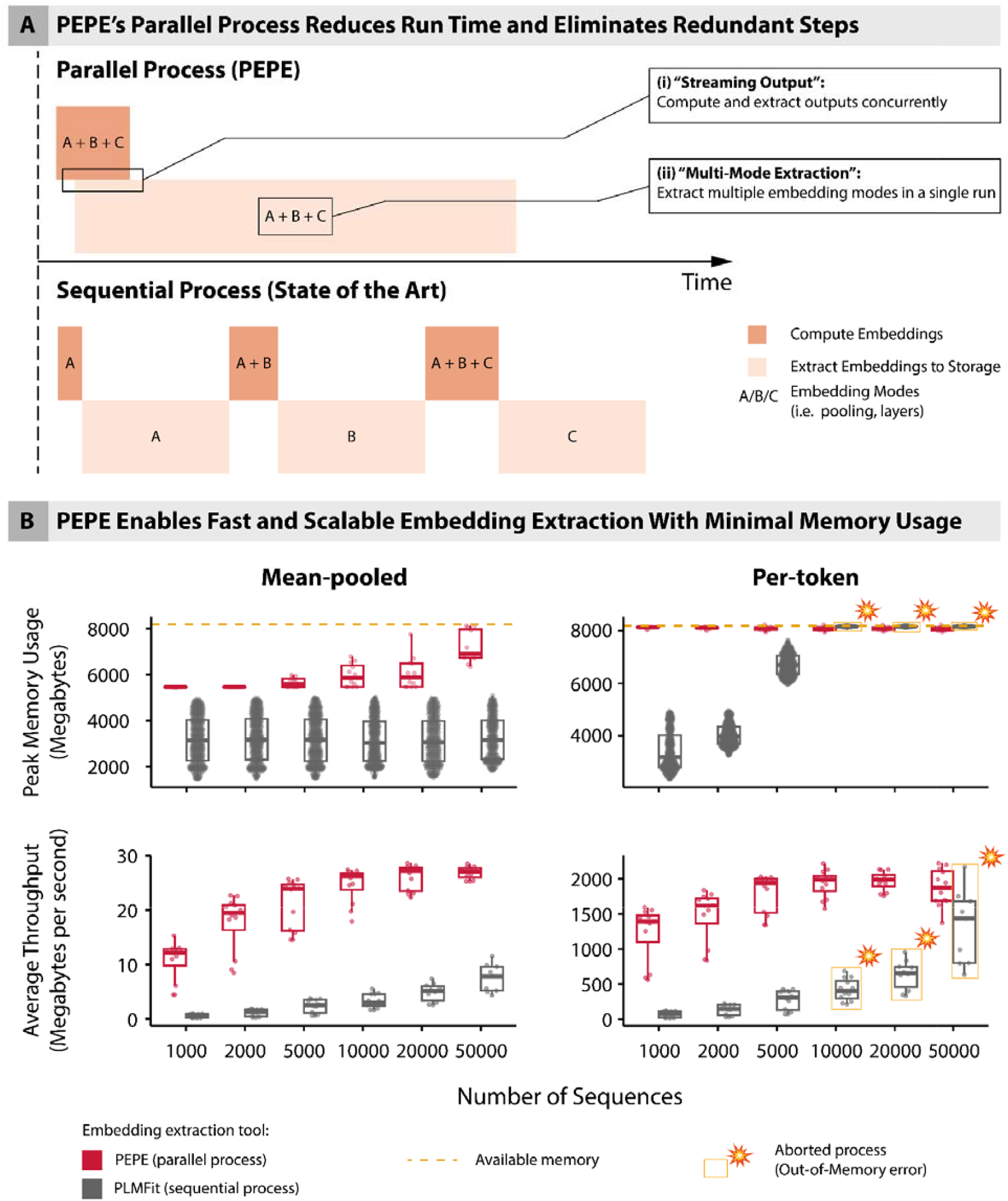
PEPE’s parallel processing enhances capacity and throughput compared to state-of-the-art methods. **(A)** Schematic comparison between parallel process used by PEPE and sequential process used by state-of-the-art (SOTA) tools such as PLMFit ^27^, AMULETY ^27,29^ and Bio-embeddings ^28^, for the extraction of embedding modes from PLMs. Orange blocks represent the computation of embeddings and beige blocks represent the extraction of one or multiple embedding modes (A/B/C) to storage. PEPE offers two key advantages: **(i)** Its “*Streaming Output*” function allows the total size of outputs to exceed the maximum available memory and reduces total run time. **(ii) *“Multi-Mode Extraction”*** further reduces run time and resource overhead by extracting all selected modes from a single embedding run. **(B)** Comparison of peak memory usage (upper panels) and average throughput (lower panels) of embedding extraction between PEPE and PLMFit (SOTA). Amino acid (AA) sequences of antibody variants (121 AAs) were collected into sets of size 1000, 2000, 5000, 10000, 20000, and 50000. Both programs were instructed to extract multiple embedding modes (per-token and mean-pooled) from all 33 embedding layers of the ESM-2 model. Per-token embeddings are the unreduced output of the embedding layers, while mean-pooled embeddings are averaged across token dimensions. All commands were run as SLURM jobs, limited to 1 CPU core, 8192 MB of CPU memory (yellow dashed line), and 1 NVIDIA A100 GPU. PEPE’s “Multi-Mode Extraction” was used to extract all 33 layers in a single run, while for the equivalent output, PLMFit’s sequential process requires a repeated run for each layer. Data points in yellow rectangles indicate aborted processes due to out-of-memory errors. All measurements were repeated 11 times (Table 3).

## 2. Results

Here, we present PEPE (**P**arallel **E**xtraction for **P**rotein **E**mbeddings), a Python command-line tool for the scalable generation of multi-modal embeddings from both public and custom PLMs, enabling embedding extraction at previously unfeasible scales (Figure 1A). We compared PEPE to PLMFit in terms of scalability and performance by using them to extract per-token and mean-pooled embeddings across all 33 layers of ESM-2 (total of 66 embedding modes per sequence) while recording memory usage over run time (see Materials and Methods). The extractions were performed with six input datasets of increasing scale (1000, 2000, 5000, 10000, 20000, and 50000 sequences of antibody variants with 121 amino acids in length). As shown in Figure 1B, although PEPE’s peak memory usage is generally higher than that of PLMFit when extracting mean-pooled embeddings, equivalent extractions had 3.6- to 18.7-fold higher throughput on average with PEPE compared to PLMFit (see Table 3). When extracting per-token embeddings that are 121 times larger than their mean-pooled equivalents, PEPE’s peak memory usage was also higher than PLMFit’s for datasets with small numbers of sequences where the average throughput increase was between 6.5- and 17.3-fold. However, when processing datasets with 10000 or more sequences, PLMFit exceeds the 8192MB memory limit, causing the process to fail due to an out-of-memory error.

In comparison, PEPE’s peak memory usage did not exceed the limit with the above-described test settings, regardless of the total output size, demonstrating its ability to scale without memory constraints with “Streaming Output” (see Methods).

Further, PEPE enables the embedding of sequences with length exceeding a model’s max length through sequence chunking and reconstruction (see Methods). An additional analogous experiment was performed where the input sequence lengths increased, but the number of sequences was kept constant (Supp. Fig. 1).

## 3. Materials and Methods

### 3.1. Program Usage

PEPE is a Python command-line tool and library for the scalable generation of multi-modal PLM embeddings from protein sequences. PEPE employs a parallel process that addresses the inefficiencies and redundancies of SOTA methods, utilizing a sequential approach. For more detailed instructions and examples, see https://github.com/csi-greifflab/pepe-cli.

A minimum of three options is required to run PEPE: a protein sequence file in FASTA format, the name of the embedding model, and a directory where outputs will be saved as Numpy arrays. The following command template extracts mean-pooled embeddings from the final layer (default options) of the specified model and stores them in the designated output directory:

~~~
pepe --model_name <model_name> \
    --fasta_path <file_path> \
    --output_path <directory_pathy> \
    --extract_embeddings <“per-token”/”mean-pooled”> \
    --layers <layer_indices>
~~~

Out of the box, PEPE supports ESM-1 ^15^ and ESM-2 family models ^2^, as well as AntiBERTa2 ^3^ and ProstT5 family models ^30^. The corresponding model strings can be found in the Github repository. There are three ways to set the embedding model using `--model_name`: (i) select a supported model by passing its model string (e.g., “esm2_t33_650M_UR50D”); (ii) select a compatible PLM by passing its Hugging Face Hub address ^31^(e.g., “alchemab/antiberta2); or (iii) select a local model by passing its directory path. Additionally, to decrease run times and conserve storage space, outputs can be saved as either 32-bit (full precision) or 16-bit (half precision) floating point numbers.

### 3.2. Handling of Long Protein Sequences

To ensure robustness against memory constraints and model-specific architectural limits (e.g., the 1022-residue limit for ESM-2), PEPE implements an automated sequence chunking and reconstruction pipeline. When the `--split_long_sequences` flag is active, the tool automatically detects the maximum sequence length permissible by the selected model’s tokenizer. Sequences exceeding this limit are partitioned into segments of length L. To preserve local biochemical context at the boundaries of these segments, users can define an overlap (stride) O using the `--split_overlap` parameter. Further, users may use the `--force_split_length` argument to manually set a maximum chunk size regardless of the model’s maximum sequence length.

After individual chunks are processed through the model’s transformer layers, PEPE reconstructs a continuous representation for the full-length protein. The tool performs a “stitching” operation that extracts the non-overlapping “payload” from each chunk. This ensures that the resulting output tensor maintains a 1:1 correspondence with the original input sequence length, eliminating redundant boundary tokens.

Further, for chunked sequences, global representations are not derived from individual chunk averages. Instead, the mean-pooled embedding is calculated across the entirely reconstructed per-token tensor. This provides a unified vector representing the full protein.

### 3.3. Embedding Modes

Multiple representations (embedding modes) of the same sequence can be extracted from a PLM (see Introduction). Embedding modes are selected by type, pooling, and layer (see Table 2). The two types of modes are conventional embeddings (outputs of embedding layers) and attention matrices (outputs from self-attention heads within embedding layers).

Currently, PEPE supports three embedding pooling options: uncompressed “per-token” embeddings, “mean-pooled” embeddings and for proteins with sub-regions known to be functionally relevant (such as the CDRs of antibodies), we also support “substring-pooling”. Here, the user supplies a CSV file with substrings that match the input sequences. Each sequence is then embedded and pooled by averaging over the substring positions.

In addition to conventional embeddings, PEPE can also extract token attention matrices. They can be extracted from individual self-attention heads of a hidden layer (per-head), averaged across all heads of a layer (per-layer) and across all heads of all layers (per-model).

Each type and pooling of embeddings can then be extracted from all or any of a PLM’s layers. The layers can be defined by supplying a list of integers with negative integers indexing layers in reverse order (−1 = final layer of PLM). PEPE can be instructed to extract embeddings from all PLM layers by passing “all” to the `--layers` option.

### 3.4. Streaming Output and Multi-Mode Extraction

PEPE’s “Streaming Output” is enabled by extracting embeddings to memory-mapped Numpy array files^32^. Memory-mapping is used to read and write to smaller parts of an array without pre-loading the entire file into memory. Here, all available memory is utilized as a buffer that is continuously filled with output data and emptied by writing it to a file. To optimize writing speed, writes are grouped into batches and executed. This allows PEPE to (i) generate output files that are larger than the system’s memory capacity and (ii) concurrently extract batches of outputs to disk as they are embedded (Figure 1A; Table 1).

**Table 1.** Comparison of embedding extraction performance between PEPE and PLMFit with increasing numbers of sequences. Values in parentheses indicate standard deviation. All computations were performed on an HPC cluster with SLURM scheduling and measurements were averaged over 11 runs. SLURM jobs were assigned one AMD EPYC 7552 CPU core, 8192 MB of CPU memory, one NVIDIA A100 GPU with CUDA version 12.8 and fast solid-state drive storage.

| Number of Sequences | Total Output Size (Megabytes) | Average Cumulative Run Time (Minutes) |  | Average Throughput (Megabits/second) |  | Fold-change |
| --- | --- | --- | --- | --- | --- | --- |
|  |  | PEPE | PLMFit | PEPE | PLMFit |  |
| Mean-Pooled Embeddings |  |  |  |  |  |  |
| 1000 | 64.5 | 1.2<br>(0.634) | 34.1<br>(30.7) | 10.7<br>(3.61) | 0.6<br>(0.333) | 18.7 |
| 2000 | 128.9 | 1.4<br>(0.585) | 34.3<br>(31.2) | 17.7<br>(5.22) | 1.1<br>(0.641) | 15.7 |
| 5000 | 322.3 | 2.7<br>(0.658) | 37.1<br>(30.9) | 21.2<br>(4.61) | 2.3<br>(1.26) | 9.1 |
| 10000 | 644.5 | 4.4<br>(0.683) | 39.1<br>(16.6) | 24.6<br>(3.23) | 3.2<br>(1.31) | 7.6 |
| 20000 | 1289.1 | 8.3<br>(0.795) | 50.6<br>(21.0) | 26.0<br>(2.37) | 4.9<br>(1.68) | 5.4 |
| 50000 | 3222.7 | 20.0<br>(0.847) | 80.3<br>(28.4) | 26.9<br>(1.12) | 7.5<br>(2.58) | 3.6 |
| Per-Token Embeddings |  |  |  |  |  |  |
| 1000 | 7927.7 | 1.2<br>(0.597) | 34.6<br>(42.1) | 1231.1<br>(395.0) | 71.1<br>(31.6) | 17.3 |
| 2000 | 15855.5 | 1.9<br>(0.649) | 34.7<br>(75.5) | 1477.1<br>(369.0) | 133.7<br>(31.4) | 11.0 |
| 5000 | 39638.7 | 3.8<br>(0.670) | 37.8<br>(143.0) | 1788.9<br>(280.0) | 273.7<br>(30.5) | 6.5 |
| 10000 | 79277.3 | 6.9<br>(0.744) | Failed | 1932.6<br>(196.0) | - | - |
| 20000 | 158554.7 | 13.5<br>(0.939) | Failed | 1965.9<br>(134.0) | - | - |
| 50000 | 396386.7 | 36.0<br>(5.47) | Failed | 1872.4<br>(267.0) | - | - |

**Table 2.**
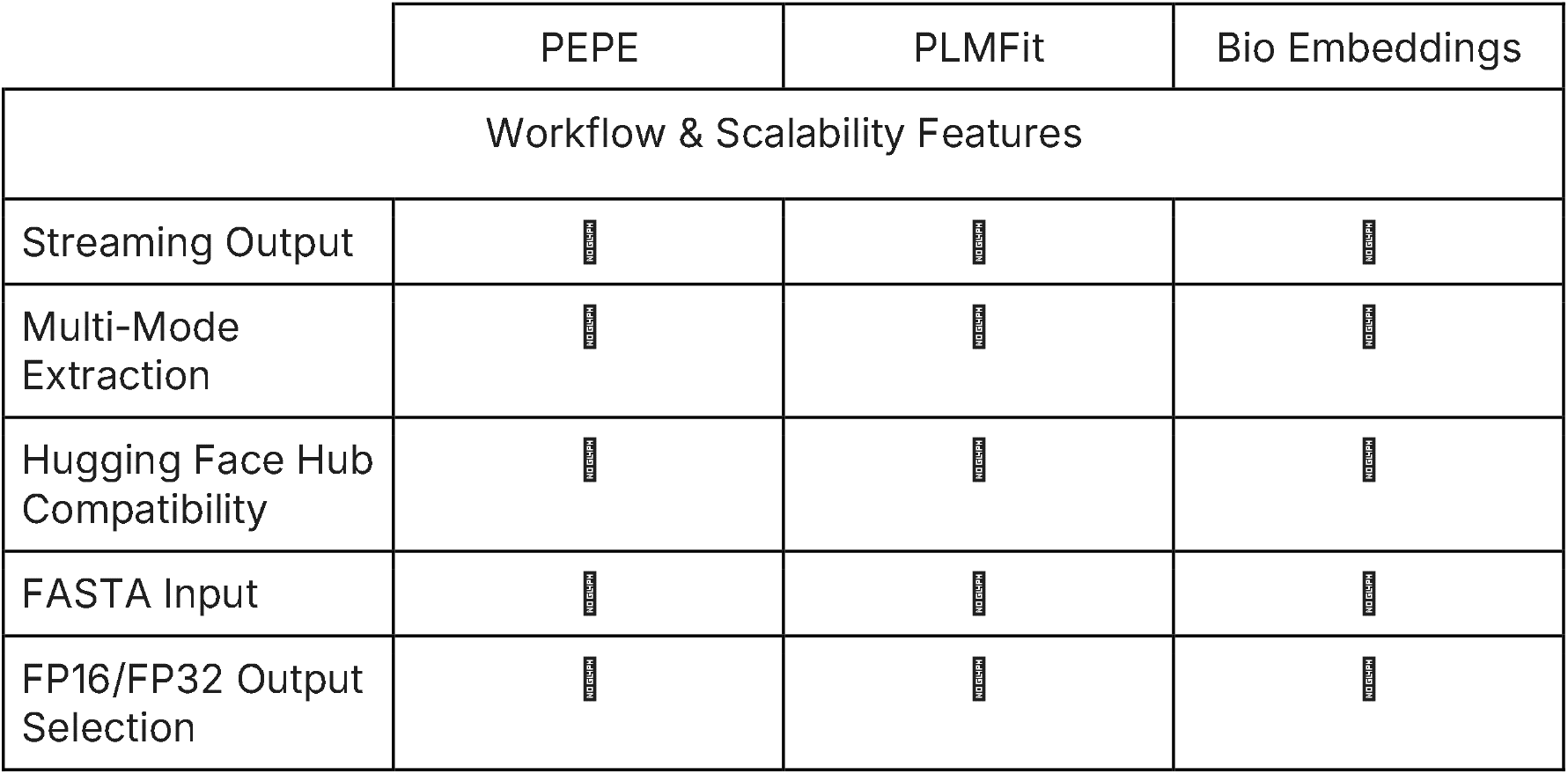
Comparison of workflow and efficiency features between PEPE and existing embedding extraction tools.

**Table 3.**
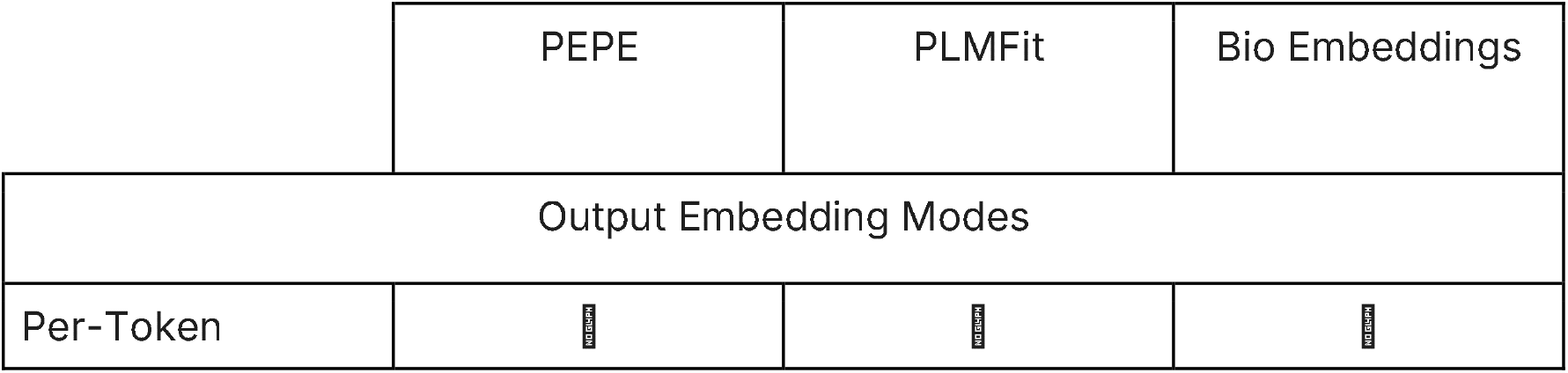
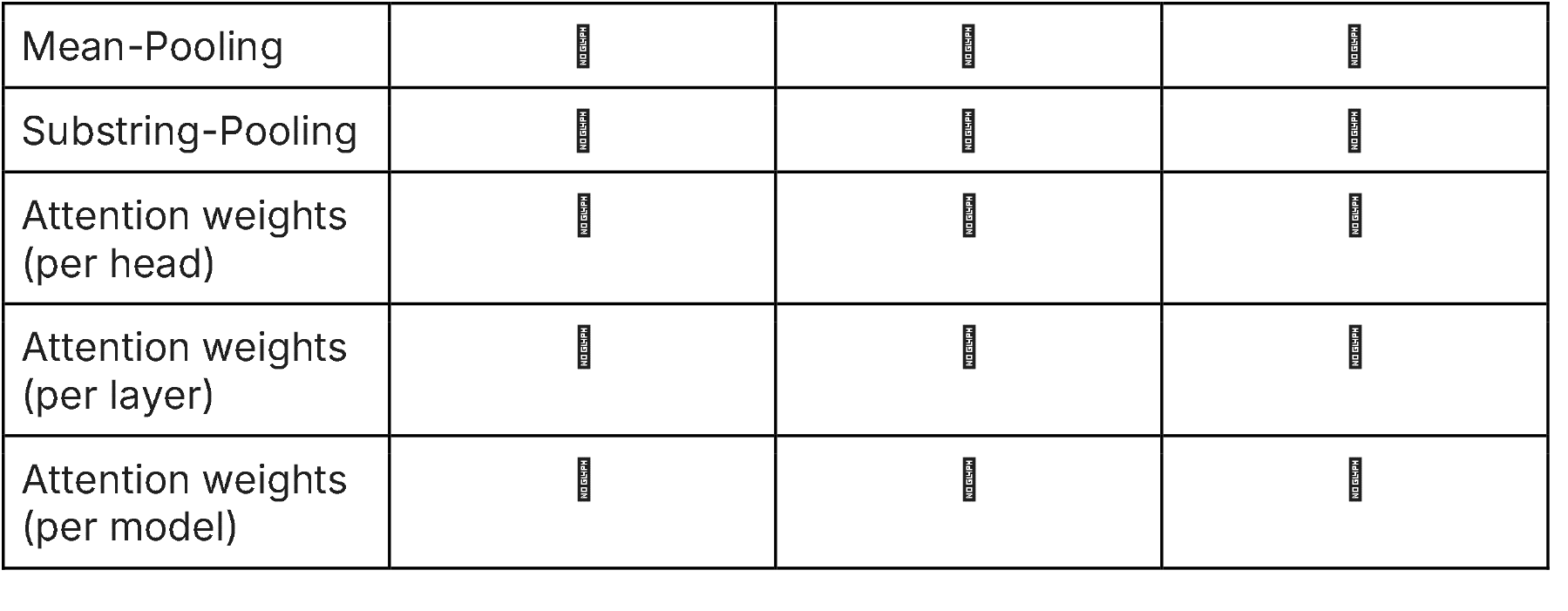
Embedding modes supported by PEPE and existing embedding extraction tools.

Embedding extraction tools using a sequential process work by embedding a given set of protein sequences, extracting a single embedding mode, typically the mean-pooled outputs of the final embedding layer. In a scenario where multiple embedding modes are extracted, the identical embedding procedure must be repeated for each additional embedding mode. PEPE avoids redundant calculations through “Multi-Mode Extraction”, where each embedding mode is assigned a separate memory-mapped file and all are extracted from a single forward pass of the PLM.

### 3.5. Comparing Scalability of Multi-Modal Embedding Extraction between PEPE and SOTA

Both PEPE and PLMFit (representative of SOTA) were installed in isolated virtual Python environments, as per the authors’ instructions. A total of 339482 antibody heavy chain variant sequences with uniform length (L = 121 amino acids) were sourced from a deep mutational scanning study ^33^. Six datasets with increasing scale (1000, 2000, 5000, 10000, 20000 and 50000 sequences) were subsampled and converted to FASTA and CSV format as input files for PEPE and PLMFit, respectively. “esm2_t33_650M_UR50D” (ESM-2), a 650 million parameter PLM from the ESM-2 family with 33 layers and 1280 dimensions (D), was used as an embedding model.

For each of the six datasets, PEPE and PLMFit were then used to extract mean-pooled (1280 dimensions) and per-token (121 * 1280 dimensions) embeddings from each of the model’s 33 layers at “half-precision” (16-bit floating point numbers). As sequential processes are typically only able to extract either mean-pooled or per-token from a single layer at a time, PLMFit required 66 separate processes per dataset (396 total) for the experiment. The equivalent extractions were performed with either two separate PEPE processes per dataset (mean-pooled and per-token) or a total of twelve. From a Python script, each run was executed as a subprocess. Using the ‘psutil’ Python library, the resident set size (RSS), or physical memory usage, of each process was recorded at 0.1-second intervals. While peak memory usage (Figure 1B, upper panels) is shown for each process, average throughput (Figure 1B, lower panels) was computed by dividing the total output size by the cumulative run time of extraction processes across all layers of a given dataset size.

To evaluate scaling with increasing sequence lengths, we generated 7 datasets of 1000 random amino acids with increasing length per sequence (50,100, 200, 500, 1000, 2000 and 5000). As shown in Supplementary Figure 1, the performance differences were similar compared to scaling number of sequences. Additionally, PLMfit is unable to embed sequences that exceed ESM-2’s maximum sequence length of 1022, whereas PEPE can handle arbitrary sequence lengths through sequence chunking and reconstruction (see Methods).

All computations were performed on an HPC cluster with SLURM scheduling ^34^ and repeated 11 times. SLURM jobs were assigned one AMD EPYC 7552 CPU core, 8192 MB of CPU memory, one NVIDIA A100 GPU with CUDA version 12.8 ^35^ and fast solid-state drive storage. In cases when outputs are written to spinning disk hard drives, we recommend increasing batch size for better performance. PEPE and PLMfit were confirmed to generate identical embeddings within rounding error margins using Numpy’s ‘allclose’ function.

## 4. Discussion

Although embeddings extracted from the final transformer layer of a PLM have been shown to be useful features for transfer learning tasks (see Introduction), determining the optimal embedding mode (e.g., embedding layer, pooling, etc.) requires extracting and comparing multiple embedding modes. The scale of an extraction is determined by the number of input sequences and the number and dimensionality of selected embedding modes (see Materials and Methods). Although PEPE only requires a single run per input dataset, mean-pooled and per-token embeddings were extracted separately to ensure comparability with PLMFit runs. Alternative embedding programs, such as AMULETY ^29^ and Bio-embeddings ^28^ were not evaluated because they only supported extracting embeddings from a PLM’s final layer without requiring custom source code modifications. Nonetheless, since they also work sequentially, we expect them to share similar memory limitations when scaling the size of outputs.

We compared the performance of PEPE against an SOTA method in multi-mode embedding extraction scenarios (Figure 1B) and found that in SOTA methods, peak memory usage increases with output size until available memory is exceeded and the process fails. In comparison, PEPE successfully finished all runs in a fraction of PLMFit’s equivalent run time, while peak memory usage remained constant even with increasing output size.

Further, while we compared multiple successive single-mode processes with PLMFit to a single multi-mode extraction with PEPE by cumulative run time, it is also conceivable to compare cumulative memory usage over peak run time instead. However, since even the smallest-scale PLMFit run had a peak memory usage above 2048MB, the cumulative memory usage over 33 embedding layers would far exceed the available memory capacity (total output size > 64GB) and would require a dedicated GPU for each process.

Finally, while SOTA methods use memory as intermediate output storage and require significant overhead to avoid out-of-memory errors, PEPE uses memory-mapped Numpy arrays that efficiently utilize all available memory as a write cache to speedup embedding extraction without exceeding limits (see Methods and Fig. 1B). Although we chose a relatively small memory capacity (8192 MB) for our experiment, reducing PEPE’s memory allocation even further would have resulted in little to no compromises in terms of run time. This is because the rate at which embeddings are generated and filled into cache far exceeds the rate at which the cache can be emptied and embeddings written to disk.

In conclusion, PEPE eliminates virtually all memory limitations inherent to SOTA methods like PLMFit, increases throughput by several fold, and enables embedding extractions at scales only constrained by capacity and write speed of the underlying storage device. As PLM capabilities, dataset sizes, and computing infrastructures continue to grow, scalable methods like PEPE will become increasingly critical for enabling future large-scale PLM-embedding-based applications.

## 8. Supplementary Figures

**Supplementary Figure 1:**
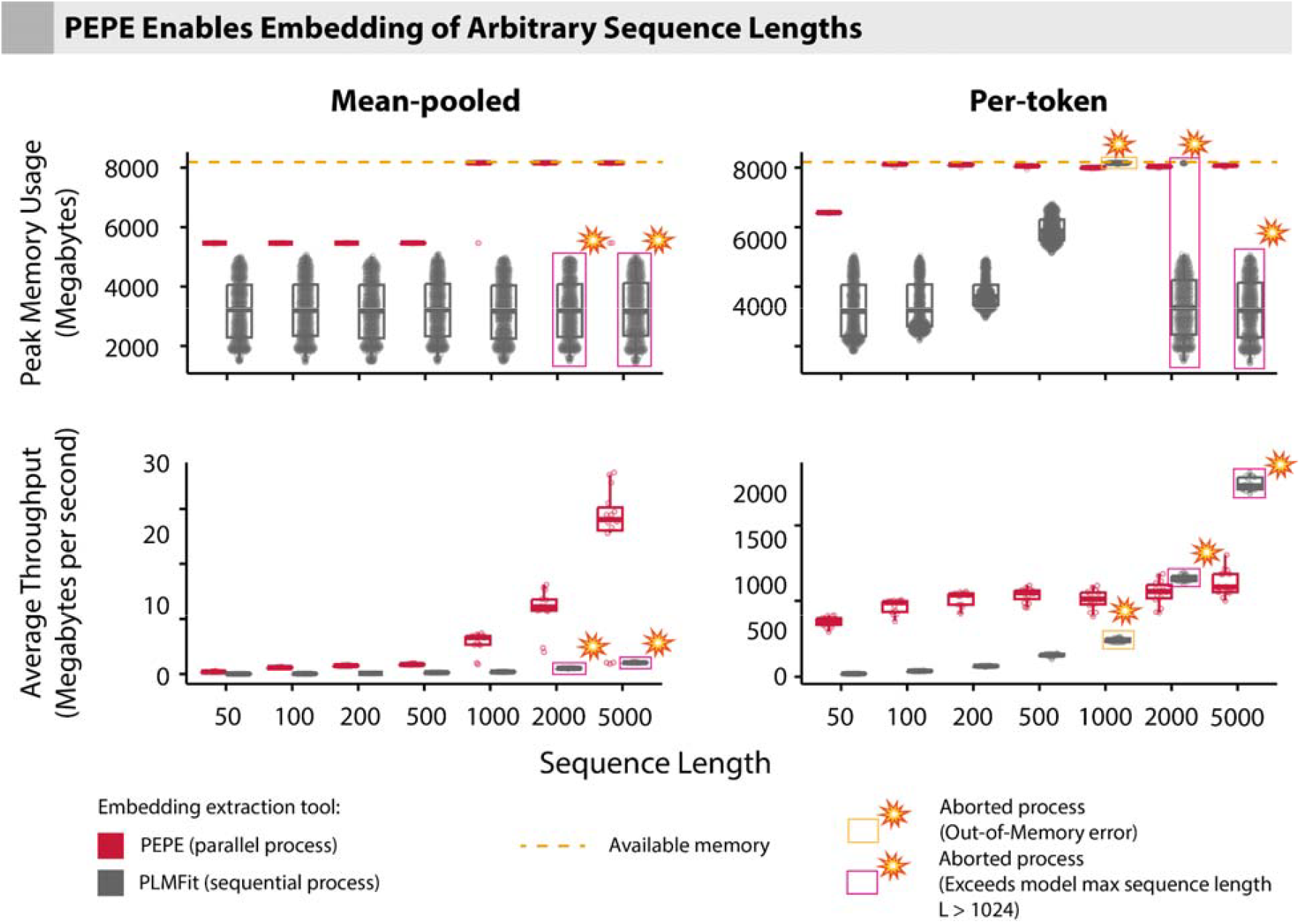
PEPE enables embedding of arbitrary sequence lengths. Comparison of peak memory usage (upper panels) and average throughput (lower panels) of embedding extraction between PEPE and PLMFit (SOTA). Sets of 1000 random amino acid (AA) sequences with sequence lengths ranging from 50 to 2000 were generated. Both programs were instructed to extract multiple embedding modes (per-token and mean-pooled) from all 33 embedding layers of the ESM-2 model. Per-token embeddings are the unreduced output of the embedding layers, while mean-pooled embeddings are averaged across token-dimensions. All commands were run as SLURM jobs and limited to 1 CPU core, 8192MB of CPU memory (yellow dashed line), and 1 NVIDIA A100 GPU. PEPE’s “Multi-Mode Extraction” was used to extract all 33 layers in a single run, while for the equivalent output, PLMFit’s sequential process requires a repeated run for each layer. Data points in yellow rectangles indicate aborted processes due to out-of-memory errors while magenta rectangles indicate aborted processes due to exceeding ESM-2’s maximum input sequence length of 1022 (plus two special flanking tokens). All measurements were repeated 15 times (Table 3).

